# EPInformer: a scalable deep learning framework for gene expression prediction by integrating promoter-enhancer sequences with multimodal epigenomic data

**DOI:** 10.1101/2024.08.01.606099

**Authors:** Jiecong Lin, Ruibang Luo, Luca Pinello

## Abstract

Transcriptional regulation, critical for cellular differentiation and adaptation to environmental changes, involves coordinated interactions among DNA sequences, regulatory proteins, and chromatin architecture. Despite extensive data from consortia like ENCODE, understanding the dynamics of cis-regulatory elements (CREs) in gene expression remains challenging. Deep learning is a powerful tool for learning gene expression and epigenomic signals from DNA sequences, exhibiting superior performance compared to conventional machine learning approaches. However, even the most advanced deep learning-based methods may fall short in capturing the regulatory effects of distal elements such as enhancers, limiting their predictive accuracy. In addition, these methods may require significant resources to train or to adapt to newly generated data. To address these challenges, we present EPInformer, a scalable deep-learning framework for predicting gene expression by integrating promoter-enhancer interactions with their sequences, epigenomic signals, and chromatin contacts. Our model outperforms existing gene expression prediction models in rigorous cross-chromosome validation, accurately recapitulates enhancer-gene interactions validated by CRISPR perturbation experiments, and identifies crucial transcription factor motifs within regulatory sequences. EPInformer is available as open-source software at https://github.com/pinellolab/EPInformer.

## Introduction

Transcriptional regulation is intricately governed by the complex interplay of DNA sequences, epigenomic signals, and three-dimensional (3D) chromatin contacts^1–3^. This process shapes gene expression and plays a crucial role in cell differentiation and environmental response. The DNA sequence interacts with various epigenetic modifications and chromatin structures to fine-tune gene expression^4^. Epigenomic signals, including DNA methylation and histone modifications, add a dynamic layer to gene regulation, influencing transcriptional activity without altering the underlying DNA sequence. Additionally, the spatial organization of chromatin, evidenced by chromatin contacts and looping, further orchestrates transcriptional regulation, bringing distant regulatory elements into proximity with gene promoters. Together, these factors constitute a multifaceted system that drives the precise and context-dependent expression of genes in living organisms.

The collaborative work and data generation efforts of consortia like ENCODE^5,6^, FANTOM^7,8^, and 4D Nucleome^9,10^ have significantly enhanced our understanding of gene regulation through epigenomics and chromatin interactions. The rich and large dataset generated by these consortia has been crucial for training powerful deep-learning methods^11^, furthering our ability to dissect and understand gene regulatory mechanisms^12–15^. These models excel by learning to predict genomic and epigenomic signals—such as transcription factor binding, chromatin contacts and accessibility, DNA methylation, and histone modifications—to improve gene expression predictions and identify regulatory elements^16–19^. This underscores deep learning’s transformative impact on computational biology and genomics^13,20,21^. However, fully understanding the complexity of *cis*-regulatory elements (CREs), such as enhancers and repressors, remains a significant challenge.

Transformer-based deep learning models have shown remarkable proficiency in predicting gene expression^22–24^. Their architecture effectively captures interactions across genomic elements, with the attention mechanism offering an advantage over traditional convolutional neural networks (CNNs) by better handling long-range genomic interactions. At the forefront of these advancements is Enformer^22^, a model excelling in predicting gene expression, protein binding, and chromatin states from DNA sequences alone. Nevertheless, Enformer’s extensive training demands limit its adaptability to unseen data from new cell types, and its efficacy in recognizing the influence of very distant enhancers remains limited (over 10 kb away from the TSS of a target gene) ^25,26^. It also does not account for three-dimensional chromatin interactions. GraphReg^27^ offers an alternative by integrating chromatin contact data to predict gene expression via a graph attention network. However, its effectiveness is constrained by the scarce availability of this data across different cell lines. Hence, there’s a pressing need for a more flexible framework for combining DNA sequences, epigenomic states, and chromatin contact data to refine predictive accuracy in cell-type-specific gene expression modeling.

To achieve this, we introduce EPInformer (a portmanteau of Enhancer-Promoter Interaction and Transformer), a scalable and efficient deep-learning framework based on the transformer architecture. Unlike other sequence-based models, EPInformer uses multi-head attention modules to directly model interactions between promoters and the potential enhancers. It integrates epigenomic signals (e.g., H3K27ac and DNase) with DNA sequences and, if available, chromatin contact data such as HiC to significantly enhance prediction accuracy. Notably, EPInformer’s streamlined architecture models gene expression in a single cell type with just 0.2% (447,149 total parameters) of Enformer’s requirements, facilitating rapid training and deployment for new cell types and reducing computational demands, a point especially important for researchers with modest computing resources. Our study rigorously tested EPInformer through a 12-fold cross-chromosome validation, confirming its superiority over existing models in predicting CAGE-seq and RNA-seq gene expression. EPInformer excels in its adaptability to various multimodal inputs. It can be trained on DNase-seq data alone or integrating DNase-seq, H3K27ac ChIP-seq, and HiC contacts for a more comprehensive analysis. Its interaction encoder effectively identifies crucial distal enhancer information, validated through CRISPR perturbation experiments. Additionally, to explore and provide interpretability of the sequence features learned by the model, we utilized TF-MoDISco-lite^29^ and TangerMEME^29^ to uncovered important transcription factor motifs within cell-type-specific enhancer sequences.

## Results

### Overview of EPInformer framework

EPInformer is a transformer-based framework for predicting gene expression by explicitly modeling promoter and enhancer interactions. The model integrates genomic sequences, epigenomic signals (e.g., DNase-seq, H3K27ac ChIP-seq), and chromatin contacts through a flexible architecture to capture their interactions. EPInformer consists of four key modules (**Fig. 1a** and **Supplementary Fig. S1**): a sequence encoder, a feature fusion layer, a promoter-enhancer interaction encoder, and a predictor module. Given a gene locus, the sequence encoder learns DNA sequence embeddings of the promoter region (1-kb sequence around the Transcription Start Site) and potential enhancers in open chromatin regions within 100kb of the TSS. Sequences shorter than 2 kb are padded with ‘N’ to reach a uniform length. Residual convolutional layers learn DNA motifs in promoter and enhancer sequences, whereas dilated convolutional layers learn motif cooperation by extracting distal sequence patterns, facilitated by the dilated convolution operator^12^. Convolutional and pooling operations in the sequence encoder work together to learn a comprehensive sequence embedding, preserving key features of the DNA sequence as shown by several past approaches^30–33^. The sequence encoder can also be pre-trained with fully connected layers to predict epigenomic signals (e.g., H3K27ac ChIP-seq) from potential enhancer regions (**Methods** and **Supplementary Fig. 1S**). This pre-training process accelerates the optimization of EPInformer and provides the model with a compositional understanding of enhancer sequence patterns before it is fully trained to predict gene expression data. Moreover, this pretrained sequence encoder enhances interpretability and helps to uncover the key motifs at the putative enhancer of the target gene.

**Fig 1.**
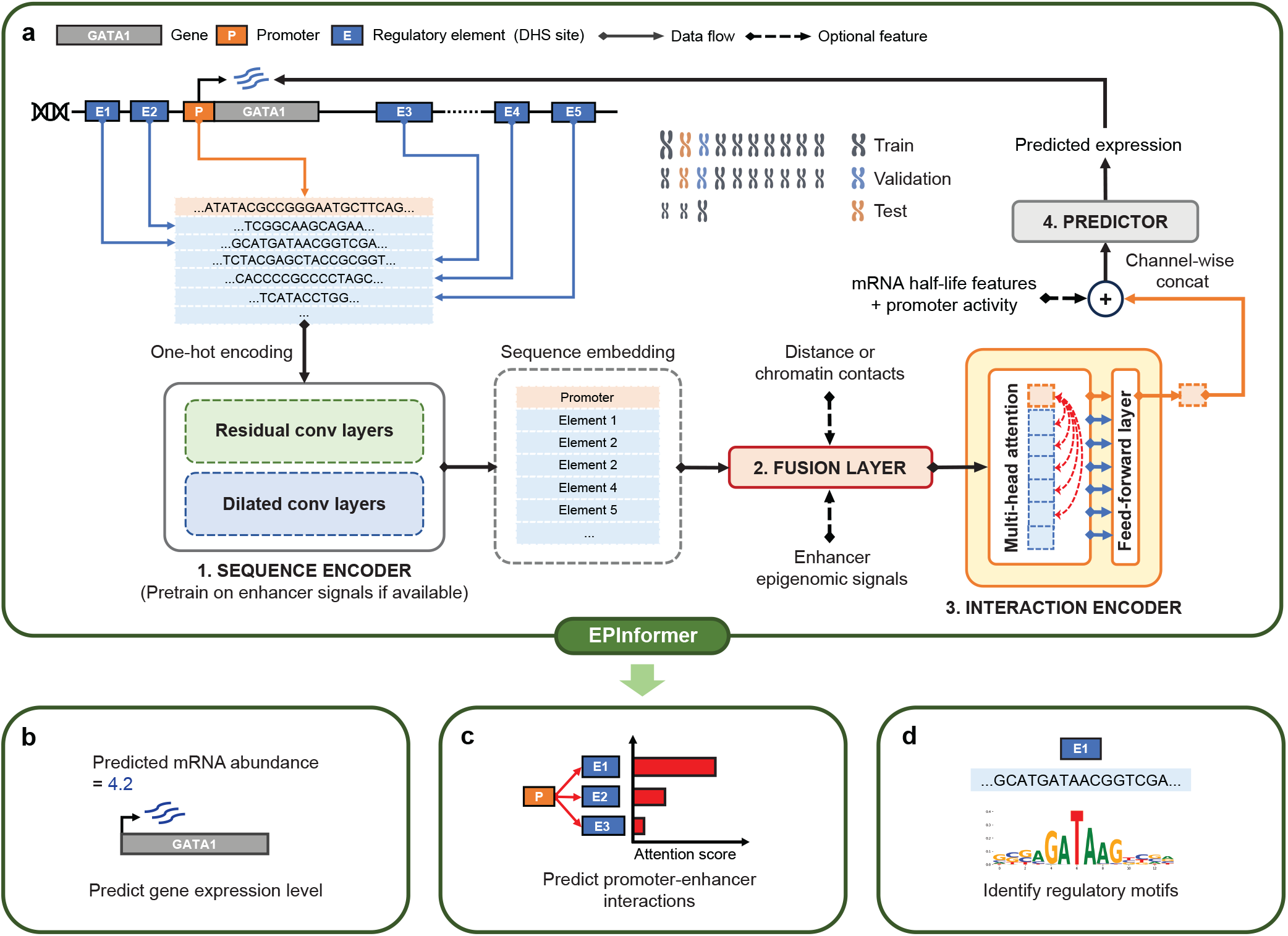
Overview of the EPInformer framework for gene expression prediction by integrating multimodal promoter-enhancer data. **a**, EPInformer is trained on multimodal epigenomic data and promoter-enhancer sequences to predict CAGE-seq or RNA-seq gene expression levels in specific cell types. It first creates embeddings for the promoter and putative enhancer sequences of a given gene using residual and dilated convolutions in the sequence encoder. This sequence encoder can be pre-trained on cell-type-specific enhancer signals to initialize the convolutional filters. The fusion layer optionally merges the sequence embeddings with distance, chromatin contacts, or epigenomic signals (e.g., H3K27ac and DNase). The interaction encoder employs a series of transformer encoders with multi-head attention modules designed to capture promoter-enhancer interactions. Finally, the prediction module integrates the resulting embeddings with mRNA half-life features and the promoter signal through fully connected layers to predict the gene expression. The EPInformer model is versatile for multiple tasks: **b**, predicting gene expression from promoter and enhancer sequences with multimodal epigenomic signals; **c**, prioritizing enhancers that may drive expression using the attention module of the interaction encoder, with scores derived from the average attention weights of the attention heads and layers; and **d**, identifying regulatory sequence features and transcription factor binding motifs at enhancers pinpointed by attention score for the target gene through the sequence encoder with downstream interpretation tools (e.g., TF-MoDISco-lite^29^ and TangerMEME^30^).

The fusion layer is designed to merge sequence embeddings with information such as distance to the target gene, epigenomic signals (e.g., H3K27ac ChIP-seq and DNase-seq), and chromatin contact data (e.g., HiC) between a promoter and candidate enhancer regions. It starts by concatenating the epigenomic signals of candidate enhancers with their sequence embeddings, followed by a 1 × 1 convolution block to refine the combined embedding dimension for the following interaction encoder. The fusion layer can integrate any number and type of genomic or epigenomic signals with sequence embedding for subsequent interaction modeling. This versatility enhances the model’s capability to incorporate diverse data types available to the users, boosting its performance and flexibility.

The interaction encoder, comprising transformer layers with multi-head attention, is designed to learn the interplay between promoters and potential enhancers. It derives a weighted sum from their embeddings, with attention weights based on fused sequence and epigenomic signal embeddings. Notably, the interaction encoder focuses solely on enhancer-promoter interactions, ignoring enhancer-enhancer interactions through attention masking. In addition, only the promoter representation after the final layer of the transformer encoder is passed, directly to the prediction module. This reduces the space of interactions to learn, focusing on promoter-enhancer interactions and increasing the computational efficiency of the model. This particular promoter representation encapsulates comprehensive relationships between a promoter and all candidate enhancers for the final predictor module, analogous to the CLS token functionality in BERT^34,35^. Subsequently, the predictor module, a feed-forward neural network, utilizes the promoter representation and genomic features like mRNA half-life^36^ and H3K27ac signals at the promoter region (500-bp around the transcription start site) to predict gene expression levels accurately. Importantly, EPInformer trained models can be combined with TF-MoDISco-lite^28^ and TangerMEME^29^ to identify transcription factor binding motifs at the putative enhancer region, incorporating the attention score of promoter-enhancer pairs to elucidate their impact on gene expression prediction.

EPInformer was trained to minimize the discrepancy between predicted and observed gene expression levels, as measured by RNA-seq or CAGE-seq using different feature sets. EPInformer excels in three key applications: 1) Accurately predicting gene expression levels using promoter-enhancer sequences, epigenomic signals, and chromatin contacts (**Fig. 1b**); 2) Efficiently identifying cell-type-specific enhancer-gene interactions, validated by CRISPR perturbation experiments (**Fig. 1c**); 3) Precisely predicting enhancer activity and identifying transcription factor binding motifs from sequences (**Fig. 1d**).

### EPInformer improves gene expression prediction by explicitly integrating promoter-enhancer interactions

To develop and evaluate EPInformer models for gene expression prediction, we initially used the ABC pipeline^37^ (**Supplementary Fig. 2S**; **Methods**) to identify candidate promoter-enhancer pairs for coding genes in two well-characterized cell lines, K562 and GM12878. In brief, we extracted promoter sequences from the 2-kb region surrounding the transcription start site (TSS) and candidate enhancer sequences from DNase I hypersensitive (DHS) sites, prioritizing up to 60 nearby enhancers per gene. This threshold covers 95% of potential regulatory elements within a 200-kb region centered on the TSS. For pre-training EPInformer’s sequence encoder, we collected H3K27ac ChIP-seq peaks from ENCODE for the K562 and GM12878 cell lines. We targeted 256 bp regions centered on the H3K27ac peak summits and included two additional 256 bp regions flanking each side with a 100 bp overlap. The ABC pipeline was used to calculate enhancer activity from DNase-seq and H3K27ac signals for these regions in both cell lines (**Methods**). To further enrich our dataset, we included the reverse complement of each sequence, retaining the same activity level, resulting in datasets of 764,888 and 536,890 samples for K562 and GM12878, respectively. Additionally, chromatin contacts of promoter and candidate enhancer pairs were obtained from KR-normalized HiC contact maps using the ABC pipeline.

Two gene expression datasets were curated for model training: protein-coding mRNA RNA-seq and Cap Analysis Gene Expression (CAGE) sequencing. For CAGE-seq, expression values were determined by aggregating read counts within 384-bp regions centered at each gene’s unique TSS, following Enformer’s protocol^22^. RNA-seq expression data were sourced from Xpresso^36^, pre-processed by the Roadmap Epigenomics Consortium. To mitigate the extreme dynamic range of gene expression across genes based on raw read count, we applied a log transformation.

To evaluate model performance under varying data availability scenarios, we evaluated several EPInformer models in predicting gene expression: EPInformer-PE takes in input promoter-enhancer sequences and the distance between the candidate enhancer and its target gene (TSS). EPInformer-PE-Activity extended this by incorporating H3K27ac and DNase signals of each enhancer element. The most comprehensive model, EPInformer-PE-Activity-HiC, in addition to including promoter-enhancer sequences and enhancer signals, can also leverage HiC contacts. To improve interpretability, the sequence encoders of EPInformer-PE-Activity and EPInformer-PE-Activity-HiC were pre-trained on a cell-type-specific enhancer activity dataset. Importantly, we also introduced a baseline model, EPInformer-promoter, which relies solely on promoter sequences.

To rigorously evaluate EPInformer models, we conducted 12-fold cross-chromosome validation across all chromosomes. In each fold, two chromosomes were designated for testing, two for validation, and the remainder for training, as proposed in previous studies^19,27^ (**Methods**). For benchmarking EPInformer against established models, including Enformer^22^, Xpresso^36^, and Seq-GraphReg^27^, we used the Pearson Correlation Coefficient (PearsonR) to compare predicted and observed gene expression levels.

Due to the substantial retraining demands of Enformer, we did not incorporate it into our 12-fold cross-validation framework. Instead, we conducted a separate hold-out test involving 1,639 genes for both Enformer and EPInformer models, using the original datasets from the Enformer study^22^. For Seq-GraphReg, which specializes in predicting gene expression on autosomal chromosomes, we referenced performance metrics from its original 10-fold cross-validation study^8^. Xpresso was evaluated alongside EPInformer within our 12-fold cross-validation.

As Seq-GraphReg and Enformer are trained to predict CAGE-seq expression, while Xpresso is developed for predicting steady-state RNA-seq expression, we trained and tested EPInformer models separately on both CAGE-seq and RNA-seq datasets to facilitate comparative analysis with these state-of-the-art models. Additionally, since the sequence encoder of EPInformer-PE-Activity(-HiC) was pre-trained on an enhancer activity dataset, we conducted the same 12-fold cross-chromosome validation covering all H3K27ac peak regions to evaluate the pre-trained sequence encoder’s efficacy in predicting enhancer activity.

In the CAGE-seq expression prediction experiments, EPInformer-PE-Activity-HiC consistently outperformed other EPInformer models and competing methods by a significant margin (**Fig. 2b**). It exceeded Enformer’s performance by 7.3% and 9.1% in Pearson correlation coefficient for K562 and GM12878, respectively, on the hold-out test set (**Fig. 2c**). EPInformer-PE-Activity, even without HiC data, achieved higher Pearson correlation coefficients of 0.841 for K562 and 0.847 for GM12878, compared to Enformer’s 0.775 for K562 and 0.774 for GM12878. In the rigorous 12-fold cross-chromosome validation on CAGE data, EPInformer-PE-Activity-HiC maintained high predictive accuracy, achieving Pearson correlations of 0.875 for K562 and 0.891 for GM12878, which are 13% and 17% higher than Seq-GraphReg, which also uses HiC to predict gene expression.

**Fig 2.**
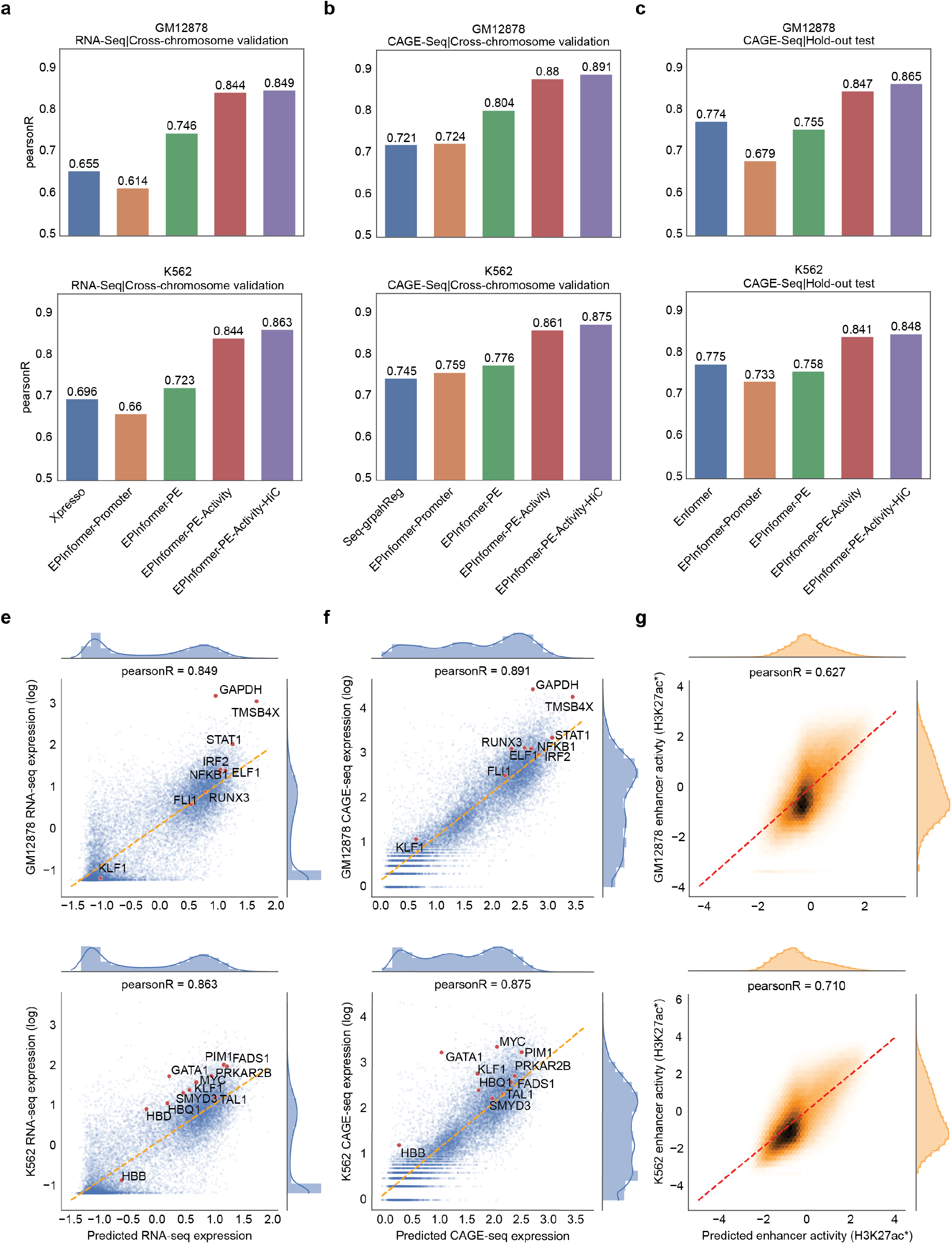
Performance of EPInformer models and baseline methods on the benchmark experiment. **a**, Comparison of four EPInformer models with Xpresso regarding their regression performance for predicting steady-state mRNA abundance, as measured by RNA-seq, in GM12878 and K562 cell lines. This evaluation was conducted using 12-fold cross-chromosome validation. **b**, Comparison of EPInformer models with Seq-GraphReg in terms of their regression performance for gene expression measured by CAGE-seq in GM12878 and K562 cell lines, utilizing 12-fold cross-chromosome validation. **c**, Comparison of multiple EPInformer models with Enformer in predicting gene expression as measured by CAGE-seq in hold-out genes within GM12878 and K562 cells. **d**, Relationship between predicted and actual RNA-seq expression levels in GM12878 (top) and K562 (bottom) cells, evaluated using 12-fold cross-chromosome validation. **e**, Relationship between predicted and actual CAGE-seq expression levels in GM12878 (top) and K562 (bottom) cells, assessed through 12-fold cross-chromosome validation. Each data point represents an individual gene. Representative cell-type-specific or housekeeping genes, activated by enhancers in K562 and GM12878 cells, are highlighted in red. **f**, The pre-trained sequence encoder quantitatively predicts enhancer activity (geometric mean of H3K27ac and DNase signals). The scatter plots show observed versus predicted enhancer activity across all DNA sequences in 12-fold cross-chromosome validation for GM12878 (top) and K562 (bottom) cells, with point density indicated by color.

In RNA-seq expression prediction experiments, three EPInformer models (EPInformer-PE, EPInformer-PE-Activity, and EPInformer-PE-Activity-HiC) outperformed Xpresso in 12-fold cross-chromosome validation for K562 and GM12878 (**Fig. 2a**). EPInformer-promoter showed lower predictive power compared to Xpresso. However, it is worth noting that this model considers a larger 20 kb region around the TSS, capturing proximal enhancer signals. In contrast, our baseline EPInformer-promoter uses only a 2 kb sequence around the TSS. EPInformer-PE, incorporating open chromatin sequences spanning 100 kb around the TSS as potential enhancers for a given gene, significantly outperformed both Xpresso and EPInformer-promoter, achieving Pearson correlations of 0.723 for K562 and 0.746 for GM12878. These findings underscore the importance of distal enhancer information for accurate gene expression prediction. Furthermore, EPInformer’s precision improved by including enhancer activity and chromatin interaction data. EPInformer-PE-Activity-HiC achieved the best performance, with Pearson correlations of 0.863 for K562 and 0.849 for GM12878.

Additionally, we conducted a 12-fold cross-chromosome validation on EPInformer’s pretrained sequence encoder, assessing its ability to predict enhancer activities (**Fig. 2g**). The strong correlation between predicted and actual enhancer activities in K562 (PearsonR = 0.71) and GM12878 (PearsonR = 0.627) cells confirms the model’s proficiency in capturing key regulatory information encoded within DNA sequences.

In summary, our EPInformer framework has demonstrated effectiveness and scalability in accurately modeling gene expression and enhancer activity. Our most sophisticated model, EPInformer-PE-Activity-HiC, significantly outperforms existing methods in predicting CAGE-seq and RNA-seq expression levels for two cell lines, providing a robust tool for predicting gene expression from sequence and multi-modal epigenomic data.

### EPInformer attention prioritizes cell-type-specific enhancers supported by long-range interaction assays

Linking candidate enhancers to their target genes through biochemical annotations remains a critical challenge in genomic research. To evaluate EPInformer’s accuracy in capturing true promoter-enhancer interactions, we focused on genes with experimentally validated enhancers. We extracted attention weights from EPInformer’s interaction encoder for these genes and compared them to known enhancer-promoter relationships. This analysis allowed us to assess whether the enhancers prioritized by EPInformer in its gene expression predictions align with experimentally confirmed regulatory interactions. The attention mechanism allows the interaction encoder to prioritize the most relevant enhancers by assigning higher weights to their sequence embeddings (Fig. 3a). We applied sum scaling normalization to the attention weights to obtain attention scores, thereby identifying and prioritizing relevant enhancers for each target gene.

**Fig. 3.**
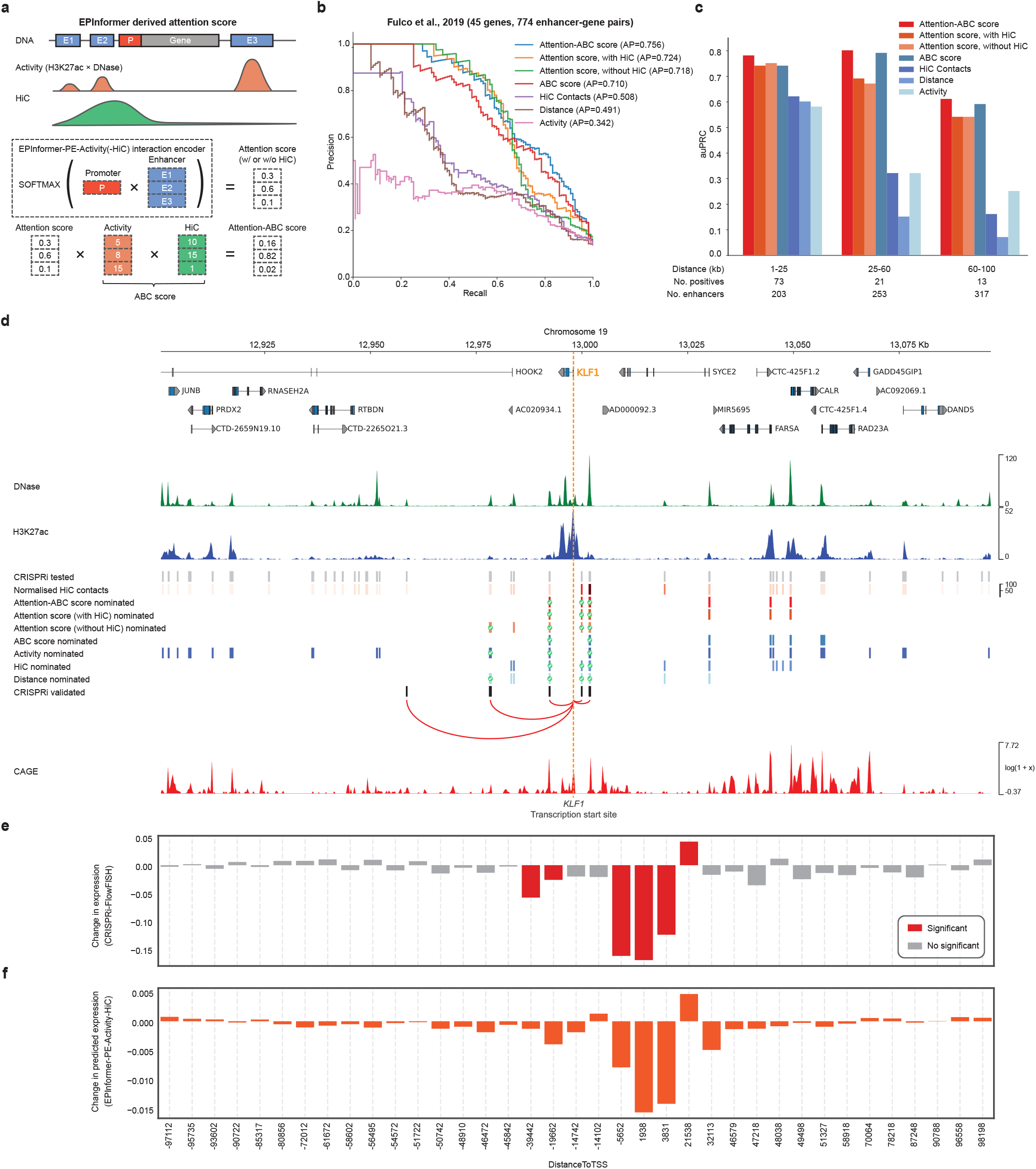
EPInformer attention prioritizes cell-type-specific enhancers corroborated by long-range interaction assays. **a**, Calculation of the attention score and attention-ABC score from EPInformer-PE-Activity-(HiC). The red and blue boxes represent three candidate enhancers (E1, E2, and E3) located near the promoter (P) of the gene (grey box). The simplified calculation of the attention weight for the promoter-enhancer pair from EPInformer-PE-Activity-(HiC) is shown in the dotted box, with red and blue boxes representing the embeddings of the promoter and candidate enhancers, respectively. The calculation of the attention-ABC score is depicted at the bottom. The values for the attention scores, DNase, H3K27ac, and HiC are expressed in arbitrary units and are not shown to scale. **b**, Precision-recall plot for classifiers of enhancer-gene (E-G) pairs. Positive E-G pairs are those for which perturbation of the candidate enhancer significantly decreases expression of the gene. The Precision-recall curves represent the performance of the attention score of EPInformer-PE-Activity-(HiC), Attention-ABC score, and other assays on classifying 737 E-G pairs of 43 genes screened by CRISPRi-FlowFISH. **c**, Area under the precision-recall curve (auPRC) represent the performance of the E-G classifiers shown in **b** on classifying enhancer-gene pairs at three different distance ranges. **d**, the nominated enhancers of all classifiers at an 85% recall threshold in **b** for *KLF1*. The locus is displayed from top to bottom as follows: candidate enhancers (light grey) defined as DNase peaks within 100 kb of the *KLF1* TSS; normalized HiC contacts between the *KLF1* TSS and the candidate enhancers. The subsequent loci present the nominated enhancers from various E-G classifiers: Attention-ABC score, the attention score of EPInformer-PE-Activity-HiC, the attention score of EPInformer-EP-Activity, ABC score, enhancer activity, HiC contact, and distance between the enhancer and *KLF1* TSS. Further down are depicted the five CRISPRi-FlowFISH validated enhancers (black box and red arc) that upon perturbation significantly decrease *KLF1* expression. The bottom red track displays the CAGE signals across the 200-kb region centered at the *KLF1* TSS. The orange dotted line marks the TSS of *KLF1*. The tick icons indicate true positive enhancers nominated by a given E-G classifier. e, Bar plot illustrating the fractional change in gene expression resulting from CRISPRi-FlowFISH perturbation on candidate regulator of *KLF1* within 100 kb of the TSS. The red bar indicates the element that caused a significant change in expression during the CRISPRi perturbation. f, Bar plot showing the fraction changes in predicted gene expression resulting from in-silico perturbations of each candidate regulator of *KLF1* using EPInformer-PE-Activity-HiC.

To validate our predictions, we used data from recent CRISPRi-FlowFISH based large-scale enhancer screens^37^. CRISPRi-FlowFISH is a powerful enhancer screening assay that integrates KRAB-dCas9 interference to target putative enhancers within a locus, coupled with RNA FISH to assess the activity of these perturbations on proximal genes. This method facilitates pooled screening and high-throughput measurement of the effects of enhancer perturbation on gene expression. Additionally, this study introduces the ABC (Activity-by-Contact) score, a computational approach designed to prioritize relevant enhancers to the target genes, which is currently recognized as one of the leading methods for this task. Despite chromatin contact data providing a strong experimental basis for enhancer-promoter interactions, this assay is not available for many cell types. Therefore, we derived two attention scores from EPInformer-PE-Activity and EPInformer-PE-Activity-HiC: the former requires only DNase-seq and H3K27ac ChIP-Seq data, while the latter requires additional HiC data. Additionally, inspired by the ABC score formulation, we developed a new score called the attention-ABC score, which is the product of the attention score of EPInformer-PE-Activity and the original ABC score.

To systematically evaluate the efficacy of attention scores to prioritize relevant enhancers for a particular gene, we collected 774 tested enhancer–gene pairs (within 100-kb to the TSS) from 45 genes in the CRISPRi-FlowFISH study performed on the K562 cell line. Note that chromosomes containing the test genes were excluded from the training process when deriving the attention scores.

As measured by the area under the precision-recall curve (AUPRC), all three attention-based scores demonstrated higher accuracy in prioritizing CRISPRi-validated enhancer-gene pairs compared to the ABC score, relative distances, Hi-C contacts, and enhancer activities (**Fig. 3b**). This is noteworthy because the ABC score relies on HiC, H3K27ac, and DNase as inputs, whereas the attention score derived from EPInformer-PE-Activity does not require HiC data. The attention score of EPInformer-PE-Activity thus can be used to prioritize enhancer-gene pairs in a broader range of cell lines that lack HiC data. As expected, we observed that the performance of the attention score could be enhanced by incorporating HiC contacts as input, achieving a higher AUPRC of 0.724 compared to the attention score of EPInformer-PE-Activity. Notably, the performance of the attention score can be significantly improved by multiplying it with the original ABC score (AUPRC = 0.756 versus 0.71).

Moreover, we compared attention scores with ABC scores in prioritizing validated enhancers at three different distances. All three attention-based scores demonstrated a higher Area Under the Precision-Recall Curve (AUPRC) than the ABC score for identifying validated enhancers within 25 kb of the target gene, with 70% located within this distance. The attention-ABC score consistently outperformed the original ABC score across all relative distances and additional metrics. Notably, even without HiC data, the attention score of EPInformer-PE-Activity surpassed both distance and enhancer activity scores in prioritizing validated enhancers across all relative distances. Our results indicate that the attention mechanism in the EPInformer interaction encoder can effectively help identify relevant and validated promoter-enhancer interactions.

To illustrate attention score capability in detecting distal enhancers for a given locus, we focused on the *KLF1* gene. We considered all its candidate enhancers within 100 kb, excluding those within 1 kb of the TSS, to ignore promoter perturbations. These enhancers (grey boxes in **Fig. 3d**) underwent evaluation via CRISPRi-FlowFISH experiments and only four enhancers were found to significantly impact *KLF1* gene expression (black boxes in **Fig. 3d**). To classify candidate enhancers as active or inactive, we selected a threshold for each score corresponding to 85% recall. The attention score of EPInformer-PE-Activity achieved the highest True Positive Rate (TPR) of 80%. Both the attention-ABC score and the attention score of EPInformer-PE-Activity-HiC achieved the second-highest TPR of 50%, significantly outperforming the ABC score, which had a TPR of 29%. Despite the attention-ABC and attention score with HiC achieving higher AUPRCs than attention score without HiC in the CRISPRi-FlowFISH dataset, they failed to identify an active enhancer with low HiC contact to the target gene. In contrast, the attention score without HiC was still able to recover this nearby enhancer with high H3K27ac signals. Overall, the attention-ABC score is the most effective strategy for prioritizing candidate enhancers in cell types used for model training. Without HiC data, the attention score of EPInformer-PE-Activity emerges as the second-best strategy for predicting active enhancers.

Furthermore, to investigate whether EPInformer-PE-Activity-HiC can also accurately predict changes in gene expression resulting from enhancer knockout, we conducted an additional *in-silico* perturbation study for the *KLF1* locus. We systematically masked 37 candidate regulatory elements within 100 kb of the KLF1 transcription start site (TSS), each previously tested by CRISPRi-FlowFISH. By comparing gene expression predictions before and after masking each element, we simulated the effects of enhancer knockouts. The predicted changes in expression showed strong agreement with the experimental CRISPRi-FlowFISH results, yielding a Pearson correlation coefficient of 0.88 (**Fig. 3e-f** and **Supplementary Fig. 3S**). Consistent with expectations, masking each of the five CRISPRi-validated enhancers led to decreased predicted KLF1 expression. Notably, our model also correctly identified a distal CRISPRi-validated repressor 21,538 bp downstream of the KLF1 TSS, predicting increased KLF1 expression upon its masking. These results underscore EPInformer’s capacity to accurately model complex regulatory interactions and its potential as a powerful tool for predicting the impact of genomic perturbations on gene expression

### EPInformer recapitulates TF motifs required for enhancer activity

Having established that the attention score of the interaction encoder effectively prioritizes relevant enhancers for a target gene, we aimed to evaluate the sequence patterns that may contribute to the functionality of these predicted enhancers. To this end, we used TF-MoDISco-lite^28^ and TangerMEME^29^ to examine the TF motifs within enhancers, based on EPInformer-PE-Activity’s sequence encoder pre-trained to predict enhancer activity from sequence.

Following TF-MoDISco-lite standard pipeline, we first applied DeepLIFT^38^ to calculate attribution scores on all 256-bp sequences centered at the H3K27ac summits of prioritize enhancer by our model. We then used TF-MoDISco-lite with default settings to identify the most relevant motifs associated with H3K27ac signals. Matching these motifs against the JASPAR 2024 CORE vertebrate non-redundant database^39^, we discovered shared TF motifs such as JUN, ELF1, ELK1, BACH1, and NYFA; K562-specific motifs including GATA1, GATA2, and GATA1-TAL1; and GM12878-specific motifs including SPI1 and FOSL1 (**Fig. 4a-b**). These results suggest that the pre-trained sequence encoder can help with model interpretation and to discover cell-type specific motifs important for enhancer activity.

**Fig 4.**
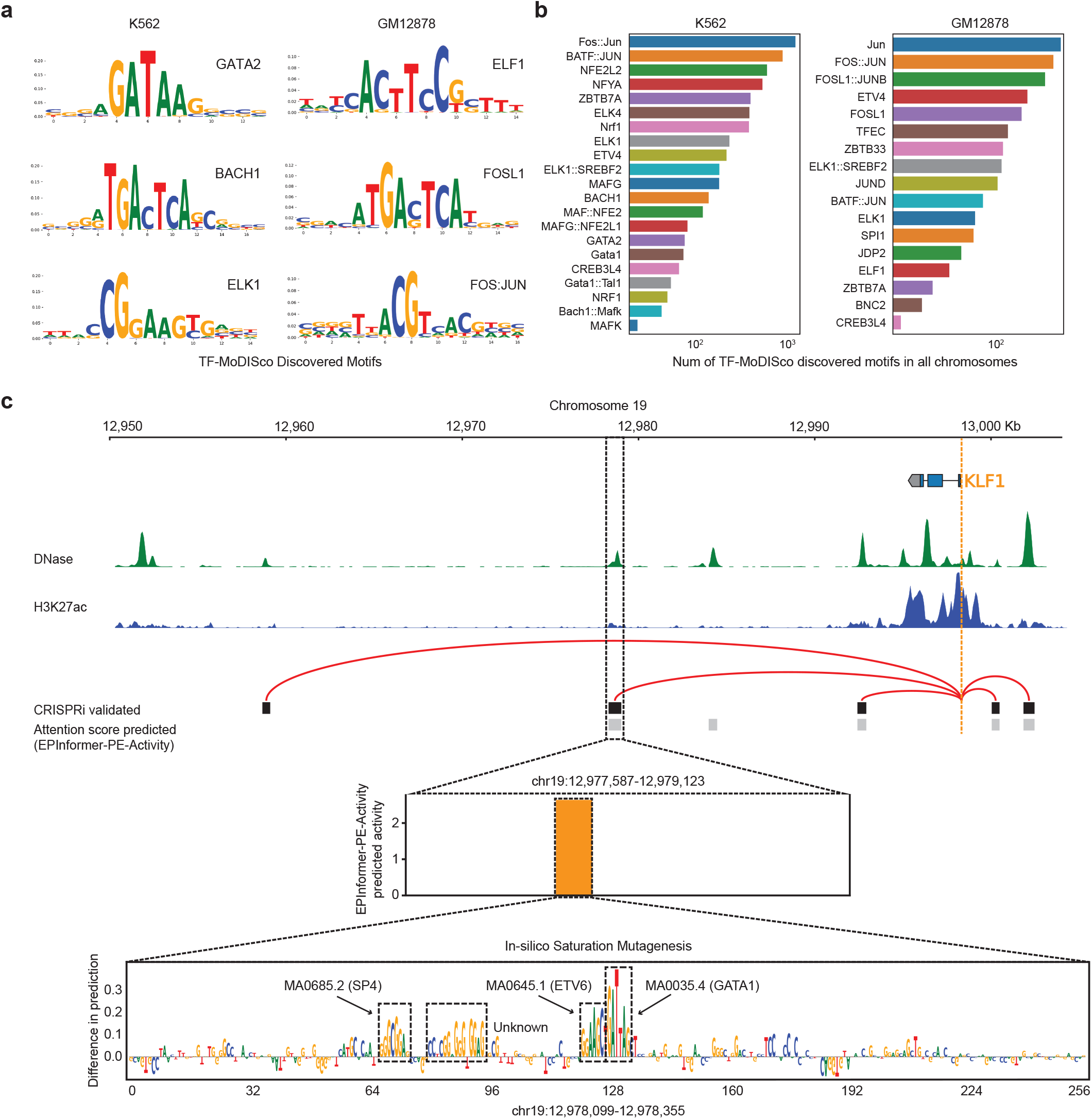
EPInformer reveals transcription factor motifs at cell-type-specific enhancers. **a**, set of three motifs enriched in K562 (left) and GM12878 (right) discovered by TF-MoDISco-lite by summarizing recurring EPInformer-PE-Activity predictive sequence patterns at all H3K27ac peaks. **b**, bar plot showing the enrichment of motifs discovered by TF-MoDISco-lite that match JASPAR 2024 CORE vertebrate non-redundant database (q-value < 0.02) in K562 (left) and GM12878 (right). **c**, EPInformer-PE-Activity predicted enhancers of KLF1 using the Attention Score, revealing several important TF motifs with its pre-trained sequence encoder on a distal enhancer (in the dotted box). The orange dotted line indicates the transcription start site (TSS) of KLF1. The black box with a red arc represents the CRISPRi-validated enhancer, while the grey box denotes the EPInformer-PE-Activity predicted enhancer. The bar plot displays the predicted enhancer activities, defined as the geometric mean of the H3K27ac and DNase signals, of 256-bp sequences tiling the putative enhancer locus (chr19:12,977,587-12,979,123). The in-silico mutagenesis (ISM) attribution score at the regions (chr19:12,978,099-12,978,355) with top predicted enhancer activity shown at the bottom. Sequences with high scores are highlighted in the dotted boxes, with labels and arrows indicating the ID and names of the resembling known TF motifs in the JASPAR 2024 CORE vertebrate non-redundant database.

Additionally, we aimed to uncover transcription motifs that may be required for KLF1 enhancers in K562 cells. Focusing on the most distal predicted enhancer at chr19:12,977,587-12,979,123, located 19,662 bp from KLF1, we first identified the window (256-bp resolution) with the highest predicted enhancer activity. Subsequently, we used TangerMEME within this window to obtain attribution scores at base resolution using the ISM approach. We observed four seqlets with high attribution scores (**Fig. 4c**), which were matched against the JASPAR 2024 CORE vertebrate non-redundant database using FIMO^40^. We found three matching motifs: GATA1, SP4, and ETV6. In K562 cells, GATA1 is a well-known master regulator necessary for erythroid differentiation, playing a crucial role in the activation and repression of various genes involved in hematopoiesis ^41,42^; ETV6 is known for its involvement in hematopoiesis and oncogenesis, contributing to the regulation of genes essential for blood cell development and differentiation^43,44^; The SP4 transcription factor (TF) is a member of the SP/KLF family of zinc finger transcription factors, and is known to bind to GC-rich promoter elements and influence the transcription of target genes^41,45^. While these insights are derived from predictive analysis, they lay the groundwork for empirical validation. Future experiments, such as motif perturbations using CRISPR deletion, base editing, or prime editing, could directly assess these motifs’ influence on gene expression.

## DISCUSSION

Identifying promoter-enhancer interactions and decoding the cis-regulatory code remains a significant challenge in gene regulation. EPInformer, a novel transformer-based framework, significantly improves gene expression prediction by modeling promoter and enhancer sequences alongside multimodal epigenomic data. EPInformer excels by holistically integrating DNA sequence, epigenomic features and chromatin contact data, offering a refined understanding of gene regulatory patterns. The CNN-based sequence encoder learns sequence patterns of promoters and enhancers, providing insight into the cis-regulatory code. The feature fusion layer integrates epigenomic signals and chromatin contacts with sequence embeddings, enhancing the prediction power and the model’s flexibility to include additional data types. The interaction encoder explicitly models promoter-enhancer interactions, while the predictor, a feed-forward neural network, harmonizes multimodal data representations to predict gene expression levels. This approach resulted in a substantial performance increase as compared to state-of-the-art tools like Enformer, GraphReg, and Xpresso and achieving Pearson correlation coefficients of 0.875 in K562 cells and 0.891 in GM12878 cells for predicting CAGE-seq expression in a 12-fold cross-chromosome validation.

EPInformer stands out from other gene expression prediction methods due to its lightweight design and versatility. Its architecture, requiring only 0.4 million parameters compared to Enformer’s 250 million, allows for faster training speeds without sacrificing efficacy. The model completes training in just one hour on a A100 GPU (**Supplementary Table 2S**), making sophisticated gene expression modeling more accessible and user-friendly for the scientific community. Importantly, the model can be trained using only DNase-seq data if necessary. However, EPInformer’s structure can easily integrate DNA sequences with multiple types of epigenomic information and chromatin interactions, enhancing its ability to predict gene expression from diverse assays like CAGE-seq and RNA-seq. This versatility ensures broad applicability and superior performance compared to models like Enformer, GraphReg, and Xpresso.

We demonstrated that EPInformer attention scores can effectively identify relevant enhancer-promoter interactions. Importantly, this approach demonstrates higher accuracy in predicting CRISPRi-validated enhancers than state-of-the-art ABC scores. Additionally, applying downstream model interpretation tools to attention score-predicted enhancers can uncover key transcription factor motifs important for cell identity.

Future enhancements to EPInformer will focus on several key areas to further improve its performance and applicability. We plan to develop more sophisticated methods for identifying and defining candidate enhancer regions, potentially incorporating additional epigenomic markers and evolutionary conservation data. Extending the model to train on and predict gene expression across multiple cell types simultaneously will improve its generalizability and ability to capture cell-type-specific regulatory mechanisms. Given the importance of CTCF in chromatin organization, we aim to integrate CTCF binding site information to better model long-range interactions and chromatin domain boundaries. Implementing relative positional encoding schemes may improve the model’s ability to capture spatial relationships between regulatory elements.

Incorporating reverse complement sequences of enhancers in the model architecture could capture additional regulatory information and improve prediction accuracy. Integrating pre-trained DNA foundation models as sequence embeddings may enhance EPInformer’s performance by leveraging large-scale genomic knowledge. Additionally, developing more comprehensive in-silico element perturbation analyses will further validate the model’s predictions and provide insights into the functional impact of specific regulatory elements. These advancements, combined with EPInformer’s current flexibility and efficiency, aim to deepen our understanding of regulatory mechanisms and their impact on gene expression and cell type identity. By leveraging CRISPR perturbation datasets and adopting a multi-task learning approach, we expect to refine EPInformer’s predictive capabilities further. Ultimately, these improvements will contribute to a more comprehensive and accurate model of gene regulation, with broad implications for both basic research and potential clinical applications.

Despite these ambitious future directions, the current iteration of EPInformer already represents a significant leap forward in gene expression prediction and enhancer-promoter interaction modeling, providing a powerful and accessible tool for researchers to unravel the complexities of gene regulation.

## Acknowledgments

We would like to acknowledge Simon Senan, Lucas Ferreira DaSilva and other people in the Pinello Lab for helpful feedback or discussions. L.P. is partially supported by 1R35HG010717-01 and MGH 2024 Research Scholar Award. R.L. was supported by Hong Kong Research Grants Council grants GRF (17113721) and TRS (T21-708 705/20-N), the Shenzhen Municipal Government General Program (JCYJ20210324134405015), the URC fund from HKU.

## Author Contribution

L.P. and R.L conceived the study. J.L., R.L., and L.P. designed the algorithms. J.L. implemented EPInformer and evaluated the benchmark results. All authors contributed to writing the manuscript.

## Methods

### Collection and pre-processing of gene expression and epigenomic data

Our study curated three types of datasets for model training and testing: enhancer-related epigenomic data, chromatin contacts, and gene expression (**Supplementary Table 3S)**. For the epigenomic data, we acquired DNase and H3K27ac bam files for all replicates of K562 and GM12878 cell lines from the ENCODE project. Following the ABC model protocol^37^ (**Supplementary Fig. 2S**), we utilized MACS2 to call peaks from the DNase-seq bam file for each cell line, considering peaks with p<0.1. We refined these to the top 150,000 regions based on read count, extended from their summits to form 500 bp candidate enhancers, and merging overlapping regions. These extended and merged peaks were defined as candidate elements in our experiments. For promoter elements, we obtained 18,377 protein-coding genes from Xpresso, excluding histone and chromosome Y genes. Following the Xpresso study, each gene’s transcription start site (TSS) was re-centered to the CAGE peak coordinates.

Promoter and putative enhancer sequences were retrieved from the hg38 reference genome. Enhancer sequences exceeding 2 kb were truncated and realigned to center on the DNase-seq peak summit. The 2 kb region surrounding the TSS was designated as the promoter, and candidate enhancers within 100 kb of the TSS, excluding the promoter region, were assigned to the target gene. This process resulted in 305,746 promoter-enhancer pairs for K562 and 325,999 pairs for GM12878.

To estimate candidate enhancer activity, we first used the ABC pipeline to compute DNase-seq and H3K27ac ChIP-seq signals from bam files by summing read counts at the candidate enhancer region. Signals from replicate experiments were averaged and quantile normalized. Based on these normalized signals, the final enhancer activity was then calculated as the geometric mean of DNase and H3K27ac signals.

We compiled a HiC dataset to estimate contacts between promoters and candidate enhancers. HiC contacts for K562 (4DNFITUOMFUQ) and GM12878 (4DNFI1UEG1HD) were obtained from the 4DN Nucleome database. Using FANC^46^, we converted the HiC data to bedpe format and applied vanilla coverage normalization at a 5 kb resolution. The ABC pipeline then computed promoter-enhancer contacts by identifying the HiC bedpe row containing the gene’s TSS and assigning contact values to enhancer-promoter pairs based on signals at the bin corresponding to the enhancer’s midpoint.

Additionally, we incorporated mRNA half-life features from Xpresso into our model, including G/C content, lengths of functional regions (5’ UTRs, ORFs, and 3’ UTRs), intron length, and exon junction density within the open reading frame.

To train and evaluate EPInformer on gene expression prediction, we curated two gene expression datasets, as measured by RNA-seq and CAGE-seq. For CAGE, expression values were determined by aggregating read counts within 384-bp regions centered at each gene’s unique TSS, as per Enformer’s protocol. RNA-Seq expression data were sourced from Xpresso’s training set, quantified by the Roadmap Epigenomics Consortium. To mitigate the right-skewed distribution of gene expression based on raw read count, we applied log-transformation.

### Model architecture

Figure 1a illustrates the model architecture, organized into four key sections: (1) a sequence encoder with 5 residual and 4 dilated convolutional layers plus a linear layer; (2) a fusion layer featuring channel-wise concatenation and 1×1 convolution blocks; (3) an interaction encoder with 3 transformer encoders, each having a 4-head self-attention module and a feed-forward layer; (4) a predictor with three dense layers for the gene expression prediction. EPInformer processes input as a one-hot encoded matrix (A = [0,0,0,1], C = [0,1,0,0], G = [0,0,1,0], T = [0,0,0,1], N = [0,0,0,0]), sized (61, 2000, 4), comprising a promoter sequence and 60 candidate enhancer sequences for predicting gene expression. Genes with fewer than 60 candidate enhancers receive padding via zero vectors to ensure uniform dimensions. The sequence encoder first learns sequence embeddings of size (61, 64) for the promoter and its candidate enhancers. The fusion layer concatenates the distances of enhancers to the TSS, enhancer activities, and promoter-enhancer chromatin contacts with sequence embeddings on a channel-wise basis. It then reshapes the concatenated matrix to a size of (61, 64) using a convolution operator. The interaction encoder then captures the interactions between promoter and the candidate enhancers using self-attention. The attention calculation is based on the matrix operation:

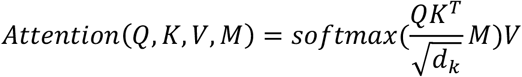

In the attention mechanism, the softmax function generates a probability distribution for promoter-enhancer interactions. A mask vector *M* is set to a value near negative infinity for padding enhancers, ensuring the interaction encoder disregards these padding embeddings. Interaction encoder learns parameter matrices 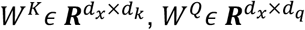 and 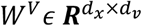 for each head, it transforms promoter-enhancer embedding 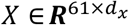 into queries *Q*_*i*_ = *X*_*i*_ × *W*^*Q*^, keys *K*_*j*_ = *X*_*j*_ × *W*^*k*^ and values *V*_*j*_ = *X*_*j*_ × *W*^*V*^. The interaction of promoter *X*_*P*_ and the *i*_*th*_ enhancer 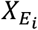 can be computed as *a_P−Ei_* = *softmax*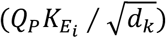, it represents the amount of weight query at promoter puts on the key at the *i*_*th*_ enhancer. Each single attention head computes its output as a weighted sum across all promoter-enhancer pairs: *a*_*P*−*E*_ × *V*. The multiple heads compute with independent parameters, and we concatenate the outputs from each head to form the final layer output, followed by a linear layer to combine them. The last transformer encoder outputs the promoter embedding with a size of 64, which embeds all promoter-enhancer pairs. Finally, the predictor concatenates the promoter embedding with 8-bit mRNA half-life features and predicts the gene expression through three dense layers.

In our enhancer activity prediction task, we engineered a model leveraging four pre-trained residual convolutional layers with filter configurations of 128, 64, 64, 128 and kernel sizes 8, 3, 3, 3. Each layer is succeeded by batch normalization, ELU nonlinearity, max pooling (size=2, stride=2), and a 1×1 convolution step. Beyond the convolutional base, the model employs two fully connected layers, each with 256 neurons, batch normalization, ReLU nonlinearity, and dropout (d=0.1). The input is a one-hot-encoded 256-bp DNA sequence aimed at predicting enhancer activities.

### Model training and evaluation

As previously proposed by Karbalayghareh et al.^27^, we implemented a 12-fold cross-chromosome validation strategy. For fold 1 to 10, chromosomes *i* and *i* + 10 were reserved for validation, while chromosomes *i* + 1 and *i* + 11 were set aside for testing. In fold 11, chromosomes 3 and 21 were used for validation, with chromosomes 22 and X allocated for testing. Fold 12 involved using chromosomes 2 and 22 for validation and chromosomes 1 and Y for testing. The remaining chromosomes were utilized for training in each fold. This evaluation procedure ensures the model is independently assessed across all human chromosomes.

All EPInformer models were implemented in PyTorch (v2.2.0)^47^ and trained on one A100 GPU with batch size of 64 using AdamW^48^ optimizer with a learning rate of 5 × 10^−4^, a weight decay of 1 × 10^−6^ and default setting for other hyperparameters: *β*_1_ = 0.9, *β*_2_ = 0.99, *ε* = 1 × 10^−8^. The models were trained using smooth L1 loss^49^ to align predictive and actual expression levels. To enhance EPInformer’s generalization and mitigate overfitting, we applied early stopping, monitoring the model’s mean square error (MSE) on the validation set and stopping training if there was no MSE improvement for six consecutive epochs. The best-performing model, marked by the lowest MSE on validation set, was retained for testing on an independent chromosome set, assessing performance through the Pearson Correlation Coefficient. For pre-training and evaluating the EPInformer’s sequence encoder, we adopted the same experimental settings as those used for EPInformer models, with the exception that this model aimed to minimize the loss between predicted and actual enhancer activity, as determined by H3K27ac ChIP-seq and DNase-seq signals (Reads per millions, RPM).

### Baseline methods

Three baseline models—Xpresso, Enformer, and Seq-GraphReg—serve as references for gene expression prediction. Enformer, a deep neural network, combines convolutional neural networks (CNNs) with transformer technology, using DNA sequences as input. It processes 196-kbp sequences to predict 5,313 genomic tracks for the human genome and 1,643 tracks for the mouse genome at 128-bp resolution. However, Enformer’s significant training requirements limit its adaptability across new cell lines, and despite its context spans around 200 kb, it can detect reliably only the impact of proximal enhancers (less than around 10 kb from the TSS)^25^. Xpresso, a deep learning model, employs CNNs to predict mRNA abundance directly from genomic sequences, focusing on promoter regions and features linked to mRNA stability within a 20 kb range of the TSS. Its reliance on proximal sequences restricts its ability to utilize information from distal enhancers. Seq-GraphReg uses graph attention networks to integrate DNA sequences and HiChIP data, predicting gene expression levels by exploiting chromatin contact signals between distal elements and promoters.

To ensure a fair comparison, we aligned the training and testing settings of EPInformer with those of Enformer. This involved using identical data splits and extracting promoter and potential enhancer sequences from the same regions Enformer was trained on. For EPInformer, gene expression values were determined by summing read counts within a 384-bp window (equivalent to three 128-bp Enformer bins) surrounding each gene’s TSS, using the same data sources (CNhs12333 for GM12878 and CNhs11250 for K562). We tested genes located in Enformer’s testing regions (1937 sequences, each 196-kbp long, covering 1639 genes) for a direct comparison.

For comparison with Xpresso, we retrained and assessed Xpresso using the same 12-fold cross-chromosome validation as EPInformer, focusing on the steady-state mRNA expression of 18,377 coding genes from the Roadmap Epigenomics Consortium. Seq-GraphReg’s performance was reported from its original study, and we presented EPInformer’s performance using an identical train-test split across all human chromosomes for direct comparison.

### Enhancer prioritization

We obtained enhancer–gene (E-G) pairs tested using the CRISPRi-FlowFISH assay from Fulco et al.^37^, which perturbs enhancers and measures gene expression changes in K562 cells (**Supplementary Table 4S**). We selected E-G pairs within 100 kb of the transcription start site (TSS), identifying 737 enhancer-promoter pairs across 43 genes. Of these, 103 were confirmed as active enhancers, showing a significant decrease in expression following CRISPRi-FlowFISH. Specifically, the active enhancers significantly reduced gene expression (adjusted p-value < 0.05) with more than 80% power to detect a 25% effect size after perturbation.

To prioritize enhancer–gene pairs with EPInformer-PE-Activity and EPInformer-PE-Activity-HiC, we extracted candidate sequences from CRISPRi-tested regions and promoter sequences based on each gene’s TSS. Enhancer activity and HiC contact data were sourced from the original study. We derived average attention weights from all heads and layers of EPInformer-PE-Activity and EPInformer-PE-Activity-HiC, excluding CRISPRi testing genes from the training set. We matched the query index at the promoter, aligning keys with different candidate enhancers. These attention weights quantified the model’s focus on each enhancer during gene expression predictions. We then normalized the attention weights for each promoter-enhancer pair, ensuring their sum equals one. These normalized attention weights were used as the attention scores to assess all enhancer-promoter pairs.

The Activity-by-Contact (ABC) score for each E-G pair was recalculated using the original code from GitHub (https://github.com/broadinstitute/ABC-Enhancer-Gene-Prediction), based on the same enhancer activity and HiC contacts used for EPInformer-PE-Activity and EPInformer-PE-Activity-HiC. The attention-ABC score was derived by multiplying the attention score of EPInformer-PE-Activity with the ABC score. Additionally, we introduced other assays to score E-G pairs, including HiC contacts, enhancer activity, and the negative distance between TSS and enhancer. The three attention-based scores were evaluated alongside the ABC score and other assays in classifying E-G pairs with significant expression changes, measured using the area under the precision-recall curve (auPRC).

### Nucleotide contribution and motif discovery

We employed TF-MoDISco-lite^28^ and TangerMEME^29^ to analyze TF motifs at putative enhancers based on EPInformer-PE-Activity’s sequence encoder, pre-trained to predict enhancer activity from sequence. TF-MoDISco-lite is a biological motif discovery algorithm which uses attribution scores from a trained deep learning model, in addition to the sequence itself, to guide motif discovery. TangerMEME is a Python package that implements the basic operations necessary to perform sophisticated genomic analyses using machine learning models. It provides a function to perform In-silico Saturation Mutagenesis (ISM) on the model given a DNA sequence of interest. ISM functions by sequentially substituting each character in a sequence with every other possible character and then assessing the change in the predictive output before and after each substitution. This observed difference is interpreted as a measure of importance or attribution, where a higher magnitude value indicates that the character change has a significant impact on the prediction, thereby suggesting its high importance. Therefore, ISM can be used to uncover block of nucleotides corresponding to TF motifs on the putative enhancer sequence.

We utilized via Captum (v0.6.0)^50^ for calculating nucleotide-specific contribution scores in sequences associated with enhancer activity. This process entailed generating 1000 dinucleotide-shuffled variants of each sequence to serve as reference points. Subsequently, the importance scores obtained from DeepLIFT^38^ for each sequence were combined with their respective one-hot-encoded matrices, yielding the final nucleotide contribution scores.

We utilized TF-MoDISco-lite v2.1.0 (available at https://github.com/jmschrei/tfmodisco-lite) to identify motifs in nucleotide contribution scores across enhancer sequences from the testing set, derived from a 12-fold cross-chromosome validation process. This tool, an efficient version of TF-MoDISco^28^, was used with its default settings to find seqlet patterns which were then compared against the JASPAR2024 CORE vertebrates non-redundant database^39^ using Tomtom^44^. To analyze nucleotide contributions to enhancer activity predictions accurately, we used *In-silico* Saturation Mutagenesis (ISM) from TangerMEME^29^ v0.2.1, generating attribution scores for each base within targeted regions.

## Data availability

We acquired epigenomic data for GM12878 and K562 cells from the ENCODE portal (https://www.encodeproject.org/), including DNase-seq (K562: ENCFF425WDA, ENCFF205FNC; GM12878: ENCFF020WZB, ENCFF729UYK), H3K27ac marks (K562: ENCFF600THN, ENCFF232RQF, ENCFF704LGA; GM12878: ENCFF269GKF, ENCFF201OHW. Additionally, CAGE data were obtained from FANTOM5 (K562: CNhs11250; GM12878: CNhs12333), and RNA-seq datasets were downloaded from the Epigenomics Roadmap Consortium (https://egg2.wustl.edu/roadmap/data/byDataType/rna/expression/57epigenomes.RPKM.pc.gz). We downloaded HiC matrices from 4DN Nucleome (https://data.4dnucleome.org/) including K562 (4DNFITUOMFUQ) and GM12878 (4DNFI1UEG1HD).

## Code availability

The code and data for EPInformer is available at https://github.com/pinellolab/EPInformer. The preprocessed training data are available at https://doi.org/10.5281/zenodo.12738705.

