## Supplementary materials for "EPInformer: a scalable deep learning framework for gene expression prediction by integrating promoter-enhancer sequences with multimodal epigenomic data"

### Supplementary tables

**Table 1S. The required data for training the gene expression prediction models.** The row highlighted in yellow are EPInformer models.

| Model | Promoter | Enhancer | DNase | H3K27ac | HiC |
|---|---|---|---|---|---|
| Xpresso | 10kb around TSS | | No | No | No |
| Seq-GraphReg | 1 Mb | | Yes | Yes | Yes |
| Enformer | 196 kb | | Yes | Yes | No |
| Promoter | 2kb around TSS | No | No | No | No |
| PE | 1kb around TSS | 100k around TSS | Yes | No | No |
| PE-Activity | 1kb around TSS | 100k around TSS | Yes | Yes | No |
| PE-Activity-HiC | 1kb around TSS | 100k around TSS | Yes | Yes | Yes |

**Table 2S. The parameters and training time of gene expression prediction models**

| Model | #Parameters | GPUs | Training time |
|---|---|---|---|
| Enformer | 250M | 64 TPU v3 cores | ~3 days |
| Seq-GraphReg | 0.3M | unknown | unknown |
| Xpresso | 0.1M | 1 Tesla A100 | ~5 mins |
| EPInformer | 0.4M | 1 Tesla A100 | ~15 mins |

### Supplementary figures

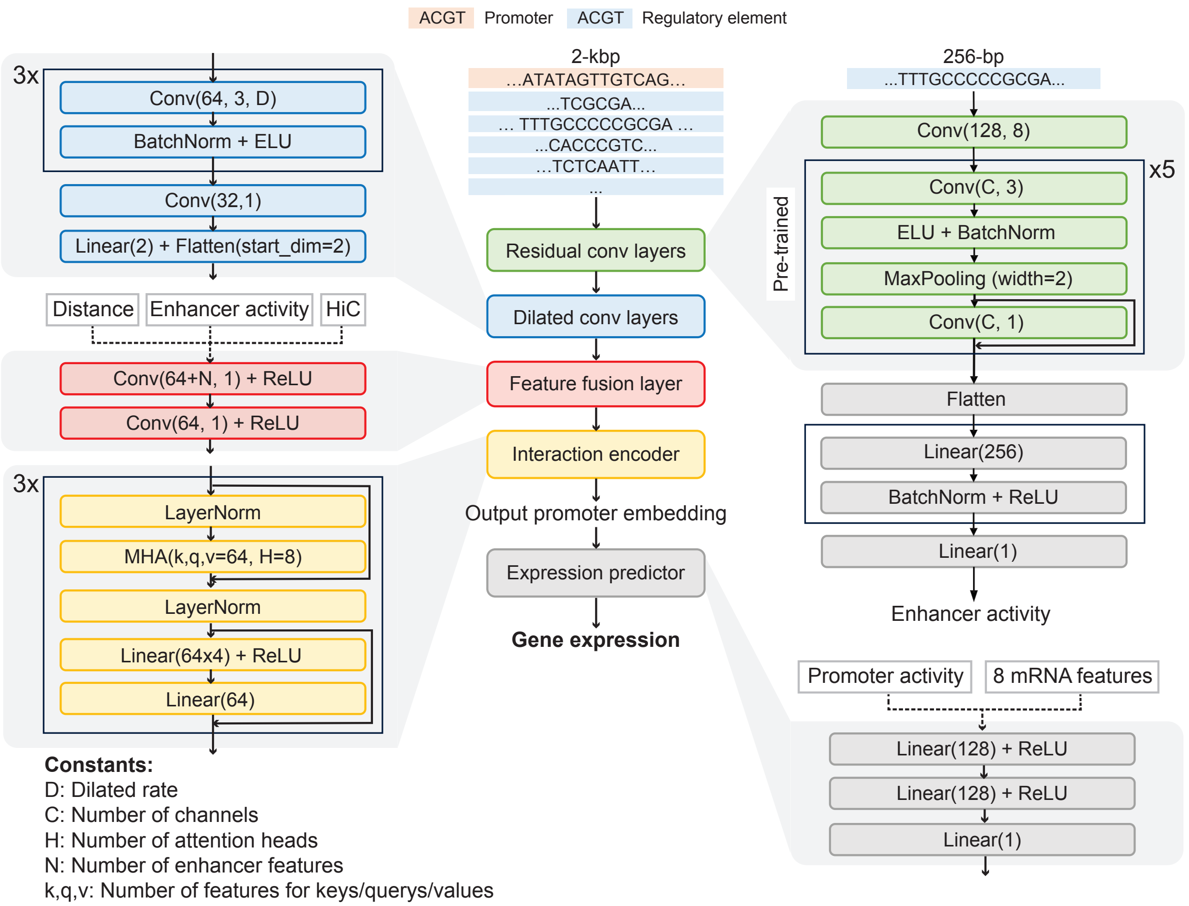


**Fig 1S. Structure of EPInformer models.** All proposed EPInformer models in our study are based on this core architecture. EPInformer-promoter masks all enhancer sequences with a zero vector and does not use any extra features; EPInformer-PE is a sequence-only model, using distance as the only additional feature. EPInformer-PE-Activity employs pre-trained residual convolution layers and takes promoter-enhancer sequences, distance, enhancer activity, promoter activity, and mRNA half-life features as inputs; EPInformer-PE-Activity-HiC includes also HiC contact data; MHA stands for Multi-Head Attention.

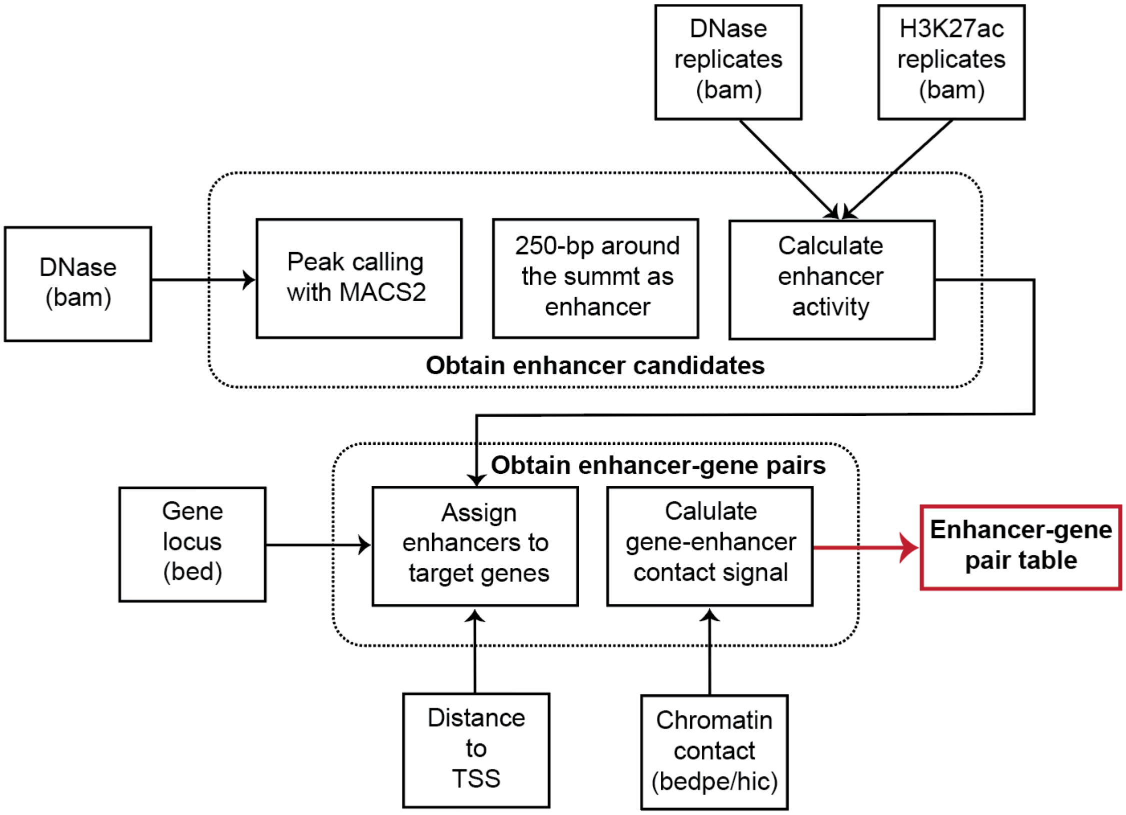


**Fig 2S. Schematic overview of ABC Pipeline for nominating enhancer-gene pairs**

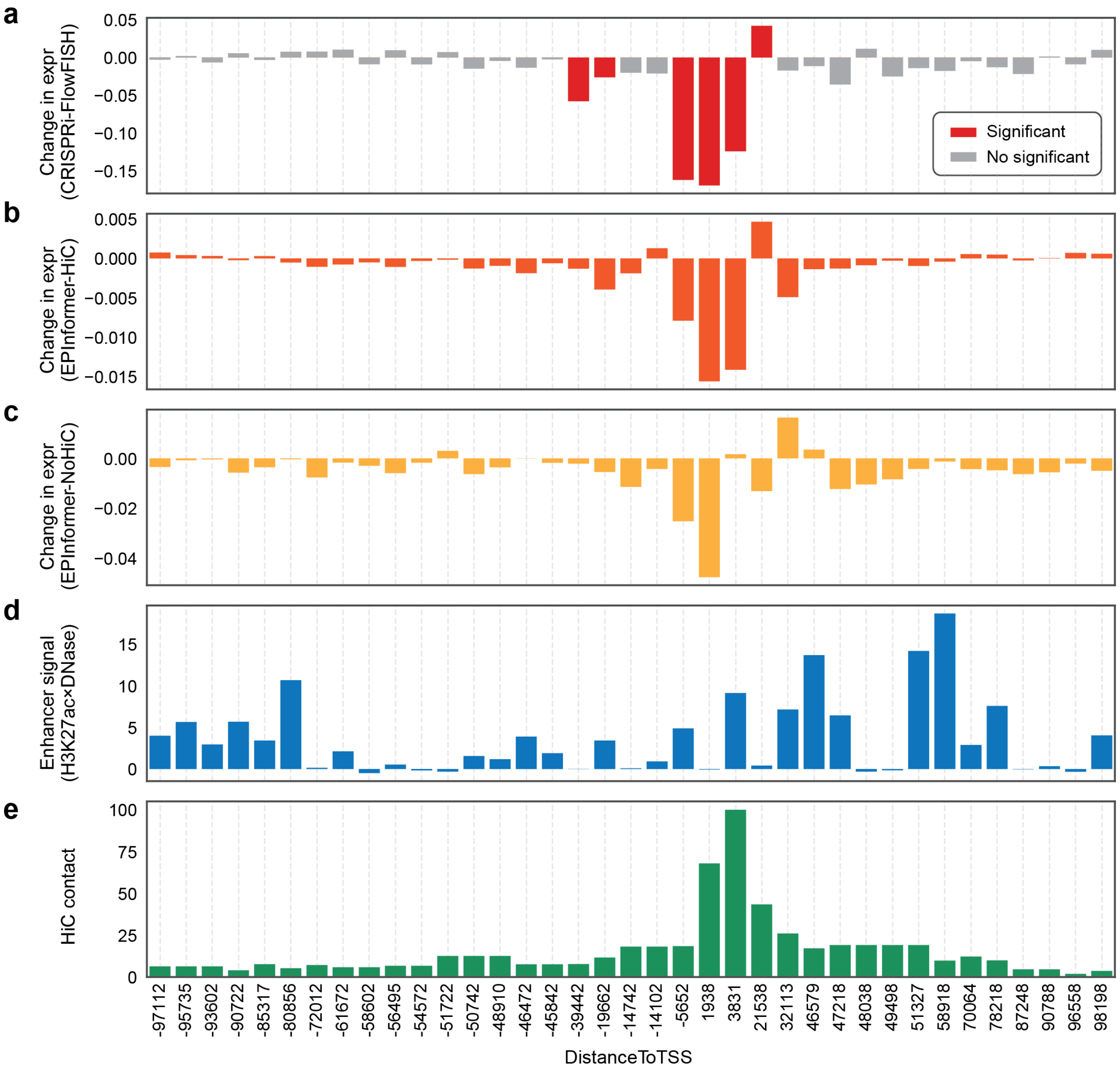


**Fig 3S. In-silico perturbation analysis of *KLF1* in K562 using EPInformer-PE-Activity-(HiC): a,** Bar plot illustrating the fractional change in gene expression resulting from CRISPRi-FlowFISH perturbation on candidate regulator of *KLF1* within 100 kb of the TSS. The red bar indicates the element that caused a significant change in expression during the CRISPRi perturbation. **b,** Bar plot showing the fraction changes in predicted gene expression resulting from in-silico perturbations of each candidate regulator of *KLF1* using EPInformer-PE-Activity-HiC. **c,** Bar plot illustrating fraction changes in predicted gene expression resulting from in-silico perturbations of each candidate regulator using EPInformer-PE-Activity. **d,** Bar plot displaying enhancer activity, defined as the geometric mean of DNase-seq and H3K27ac RPM (Reads Per Million), for each candidate regulator of *KLF1*. **e,** Bar plot indicating HiC contacts between candidate regulator and the transcription start site of *KLF1*.
